# Brain regions modulated during covert visual attention in the macaque

**DOI:** 10.1101/340703

**Authors:** Amarender R. Bogadhi, Anil Bollimunta, David A. Leopold, Richard J. Krauzlis

**Affiliations:** Laboratory of Sensorimotor Research, National Eye Institute, National Institutes of Health; Laboratory of Neuropsychology, National Institute of Mental Health, National Institutes of Health; Neurophysiology Imaging Facility, National Institute of Mental Health, National Institute of Neurological Disorders and Stroke, National Eye Institute, National Institutes of Health

## Abstract

Neurophysiology studies of covert visual attention in monkeys have emphasized the modulation of sensory neural responses in the visual cortex. At the same time, electrophysiological correlates of attention have been reported in other cortical and subcortical structures, and recent fMRI studies have identified regions across the brain modulated by attention. Here we used fMRI in two monkeys performing covert attention tasks to reproduce and extend these findings in order to help establish a more complete list of brain structures involved in the control of attention. As expected from previous studies, we found attention-related modulation in frontal, parietal and visual cortical areas as well as the superior colliculus and pulvinar. We also found significant attention-related modulation in cortical regions not traditionally linked to attention – mid-STS areas (anterior FST and parts of IPa, PGa, TPO), as well as the caudate nucleus. A control experiment using a second-order orientation stimulus showed that the observed modulation in a subset of these mid-STS areas did not depend on visual motion. These results identify the mid-STS areas (anterior FST and parts of IPa, PGa, TPO) and caudate nucleus as potentially important brain regions in the control of covert visual attention in monkeys.

---

Neuronal correlates of covert visual attention have been demonstrated in several visual cortical areas^1–7^, frontal areas^8–12^, parietal areas^13,14^ and sub-cortical regions^15–19^ of non-human primates. Causal contributions to covert spatial attention have also been demonstrated for some cortical (FEF, LIP) and sub-cortical regions (SC and pulvinar)^20–25^.

One explanation for attention-related improvements in performance is that sensory processing of the stimulus is enhanced through top-down modulation^26,27^. The strongest evidence for this idea comes from recordings in visual cortex combined with causal manipulations in FEF or pulvinar^25,28,29^. However, improvements in performance can also be achieved by other operations that do not change local sensitivity, such as changes in choice bias^30^, spatial weighting of sensory signals^23,31^, enhancing cortical communication^18^, and filtering out distractors^8,10^. The implementation of these other components involves brain areas and circuit mechanisms that are only partly understood but likely to play a central role in the control of selective attention^30–33^.

The broad coverage of fMRI offers a means to investigate attention-related modulation throughout the brain. Recent fMRI studies in monkeys have identified several cortical and sub-cortical areas modulated during covert visual attention tasks^34–38^. Most of these studies used attention to static stimuli^34,35,37,38^, with the exception of one study that used a visual motion stimulus^36^. The present study had two principal objectives. First, we aimed to replicate the findings from previous fMRI studies in monkeys^34–38^ using a different attention task paradigm. Second, we sought to identify additional attention-related brain regions that were not reported in these earlier studies, perhaps due to differences in the scanning methodology or the attention paradigm. Toward this end, we collected 62208 volumes of imaging data in two monkeys while they performed a covert attention task involving visual motion stimuli. The task was somewhat different from that used in the previous macaque fMRI studies of attention^35,36^, though was directly comparable to that used in a previous human fMRI study to identify the brain areas that show attention-related modulation to visual motion^39^.

Consistent with previous findings, we found attention-related BOLD modulation in several early visual cortical areas (V1, V2v, MT), frontal and parietal areas (FEF, vlPFC, LIP) and subcortical structures (SC, pulvinar). Importantly, we also identified significant attention-related modulation in brain areas not traditionally linked to attention – namely, mid-STS cortical areas (anterior FST and parts of IPa, PGa, TPO) and the caudate nucleus in the basal ganglia. These results identify a more complete list of cortical and subcortical brain regions that are actively recruited during covert selective visual attention in monkeys and that may contribute to different components of attention-related processing.

## Results

### Performance in the attention tasks involving visual motion

Both monkeys reliably performed three covert attention tasks (Baseline, Ignore and Attend) presented as a block design in the vertical scanner (Fig. 1; See Methods). In the Baseline task, monkeys detected luminance change at fixation. The Ignore task was similar to Baseline but included peripheral motion stimuli as distractors. In the Attend task, monkeys detected a motion-direction change in the peripheral motion stimuli. In Baseline trials (Fig. 1B), the hit rates (mean: 74%, 68%) of the two monkeys were significantly higher than false alarms on catch trials (mean: 9%, 17%) during all sessions (Chi-square proportion test; p<0.0001), indicating that monkeys based their choices on the luminance change at fixation. In Ignore trials, foil false alarms – defined as joystick releases to motion-change in Ignore trials – were significantly lower for both monkeys (mean: 6%, 9%) compared to the hit rates (mean: 72%, 68%) (Chi-square proportion test; p<0.0001), indicating that monkeys actively ignored the motion stimuli during the Ignore task. Hit rates for left (mean: 75%, 73%) and right stimuli (76%, 69%) in Attend trials for both monkeys (Fig. 1F) were significantly higher than the foil false alarms for left (mean: 6.5%, 9%) and right stimuli (5.5%, 9%) in Ignore trials (Chi-square proportion test; p<0.0001) during all sessions, showing that the monkeys successfully based their choices on either the peripheral motion stimuli (Attend task) or the central stimulus (Ignore task), even though the peripheral visual stimuli were the same between the two tasks.

**Figure 1.**
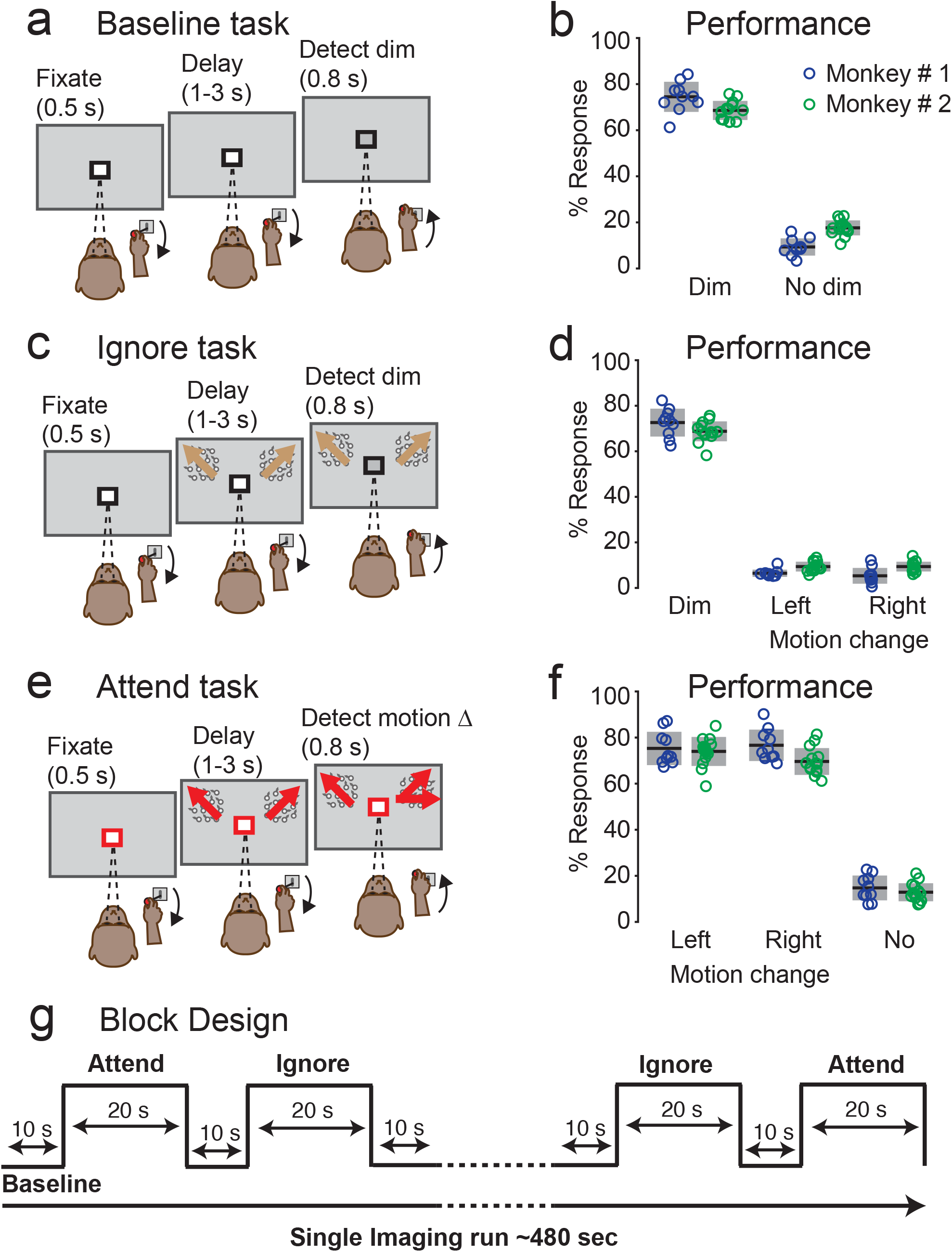
Behavioral tasks and performance. The color of the square around the central fixation spot instructed the monkey to either monitor the fixation stimulus (black in baseline (a) and Ignore (b) tasks) or the peripheral motion stimuli (red in Attend (c) task). a) Baseline task. Following 500ms of fixation, the central fixation spot dimmed on 65% of the trials during the delay period. Monkey released the joystick within 0.75 seconds of the dim to get a juice reward. b) Performance of both monkeys (Monkey # 1: blue; Monkey # 2: green) on change trials (dim) and catch trials (no dim) in the Baseline task. Circles indicate % response in each session. Horizontal black lines with gray bars indicate mean and standard deviation respectively. c) Ignore task. Following 500ms of fixation, two circular patches of random dot motion stimuli (6^0^ in diameter) appeared on either side of fixation at 8^0^ eccentricity (radius) and 10^0^ above horizontal (azimuth). During the delay period, the central fixation spot dimmed on 65% of the trials and monkey released the joystick within 0.75 seconds of the dim to get a juice reward. During the same delay period, independently of the fixation spot dimming, one of the motion stimuli changed direction on 65% of the trials. Monkey had to ignore the motion-change and hold the joystick down. d) Performance in the Ignore task. Color and symbol conventions same as (b). e) Attend task. Following 500ms of fixation, two circular patches of random dot motion stimuli appeared at the same location as in Ignore task. During the delay period, one of the motion stimuli changed direction on 65% of the trials and monkey released the joystick within 0.75 seconds of the motion-change event to get a juice reward. f) Performance in the Attend task to left and right motion-changes as well as no changes. Color and symbol conventions same as (b). g) Block Design: All three tasks were presented in a block design and the duration of each task in the block design is shown in d. Each run started with the Baseline task and was interleaved with Ignore and Attend tasks.

Overall, across 24 imaging sessions in the two monkeys we obtained 5184 blocks of trials for the attention tasks (1056 blocks each for both the Ignore and Attend tasks in monkey #1, 1536 blocks each for monkey #2). In the following, we first present the results as activation maps for the cerebral cortex, and then describe the regions of interest with attention-related modulation that we identified in both cortical and subcortical brain regions.

### Cortical activation maps during the Ignore and Attend tasks

We started by obtaining separate activation maps for the Ignore and Attend tasks. We measured the activations during the Ignore task as t-scores contrasting Ignore and Baseline and overlaid these values onto the partially inflated cortical surfaces of D99 anatomical in monkey #1’s native space for both hemispheres, along with the anatomical borders of cortical areas (Fig. 2a, 2b). We found activations in areas of the visual cortex and superior temporal sulcus (STS). Based on separate retinotopic mapping (see Supplementary Fig. S1 and Supplementary Methods), we determined that the activated voxels were predominantly in regions that mapped onto peripheral locations, consistent with them being stimulated by the moving dot stimuli.

**Figure 2.**
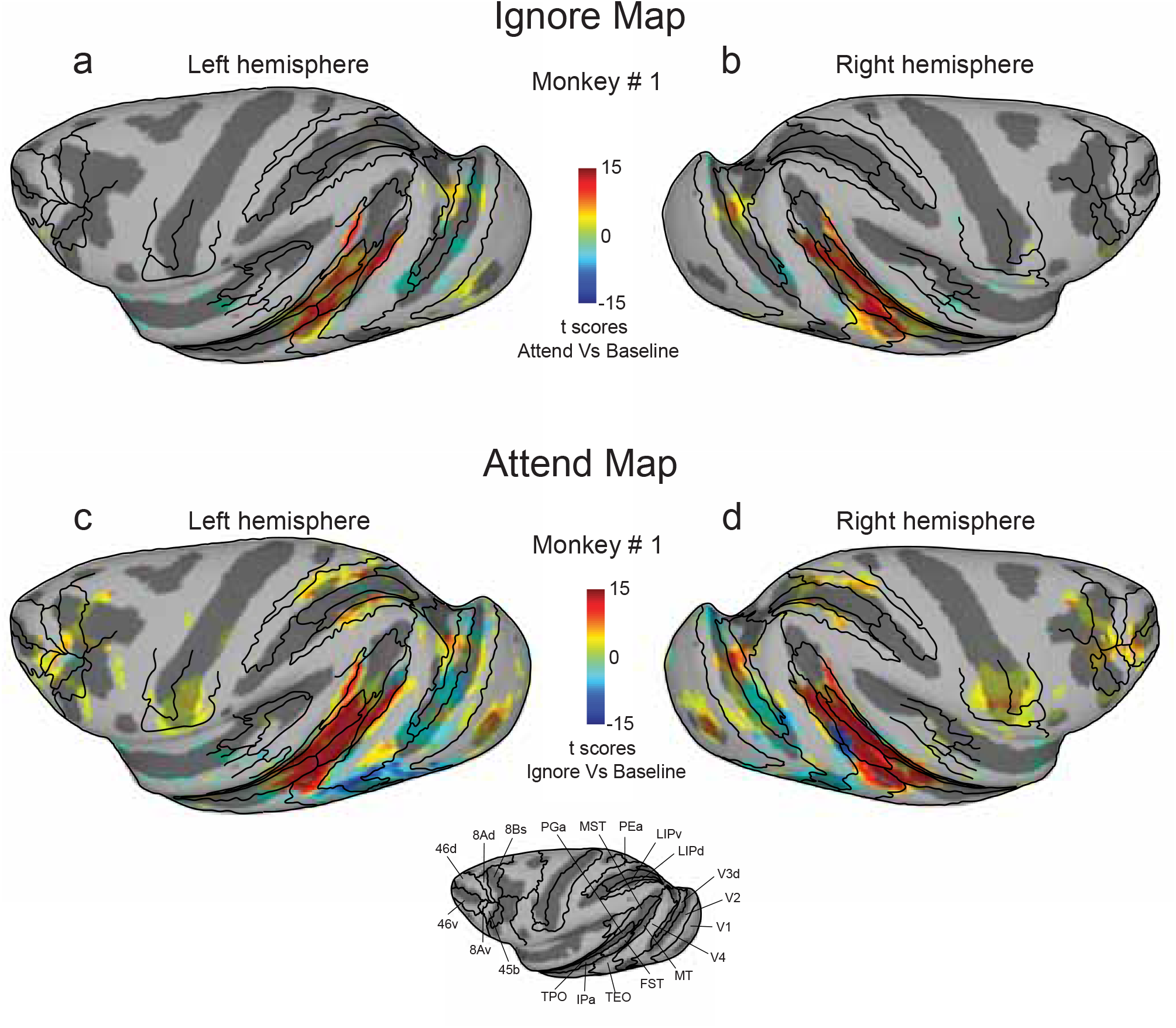
Activations during Ignore and Attend tasks (a, b) T-scores contrasting Ignore and baseline tasks show activations during Ignore task in left (a) and right (b) hemispheres of monkey # 1. (c, d) T-scores contrasting Attend and baseline tasks show activations during Attend task in left (c) and right (d) hemispheres of monkey # 1. Anatomical boundaries are labeled for the left hemisphere. T-scores were corrected for multiple comparisons (Bonferroni correction; p < 0.05, |t-score| > 5.02).

In similar fashion, we measured the activations during the Attend task as t-scores contrasting Attend and Baseline tasks (Fig. 2c, 2d), and again found activations in visual cortex and STS, and also in the intra-parietal sulcus (IPS), central sulcus (CS), arcuate sulcus (AS) and principal sulcus (PS). The positive activations predominantly corresponded to voxels that mapped onto peripheral stimulus locations. Now, in addition, there were pronounced negative activations corresponded to voxels that mapped onto foveal stimulus locations (central 2^0^ radius). One possibility is that these deactivations in foveal voxels stemmed from differences in the behavioral relevance of the foveal stimulus during the Baseline verus Attend tasks, though we cannot exclude that the deactivations arised from stimulus differences at the fovea due to the color (black versus red) or luminance changes (present versus absent) in the small fixation cue. Hereafter, we focus on the voxels corresponding to the position of the peripheral stimuli.

### Overview of attention-related modulation in cortex

To determine areas of the brain modulated during covert visual attention, we examined the contrast between the Attend and Ignore conditions, excluding voxels that mapped onto foveal locations. The resulting t-score maps (Bonferroni corrected; p < 0.05, t-score > 5.02) projected onto partially inflated cortical surfaces for both monkeys are shown in Fig. 3; the unthresholded t-score maps contrasting Attend and Ignore conditions for both monkeys including all cortical voxels are provided in Supplementary Fig. S2. The t-score maps provide a broad overview of the cortical areas that showed consistent attention-related modulation: areas of the visual cortex, superior temporal sulcus (STS), intra-parietal sulcus (IPS), central sulcus (CS), arcuate sulcus (AS) and principal sulcus (PS). We also observed significant attention-related modulation in several subcortical regions that we describe in a later section.

**Figure 3.**
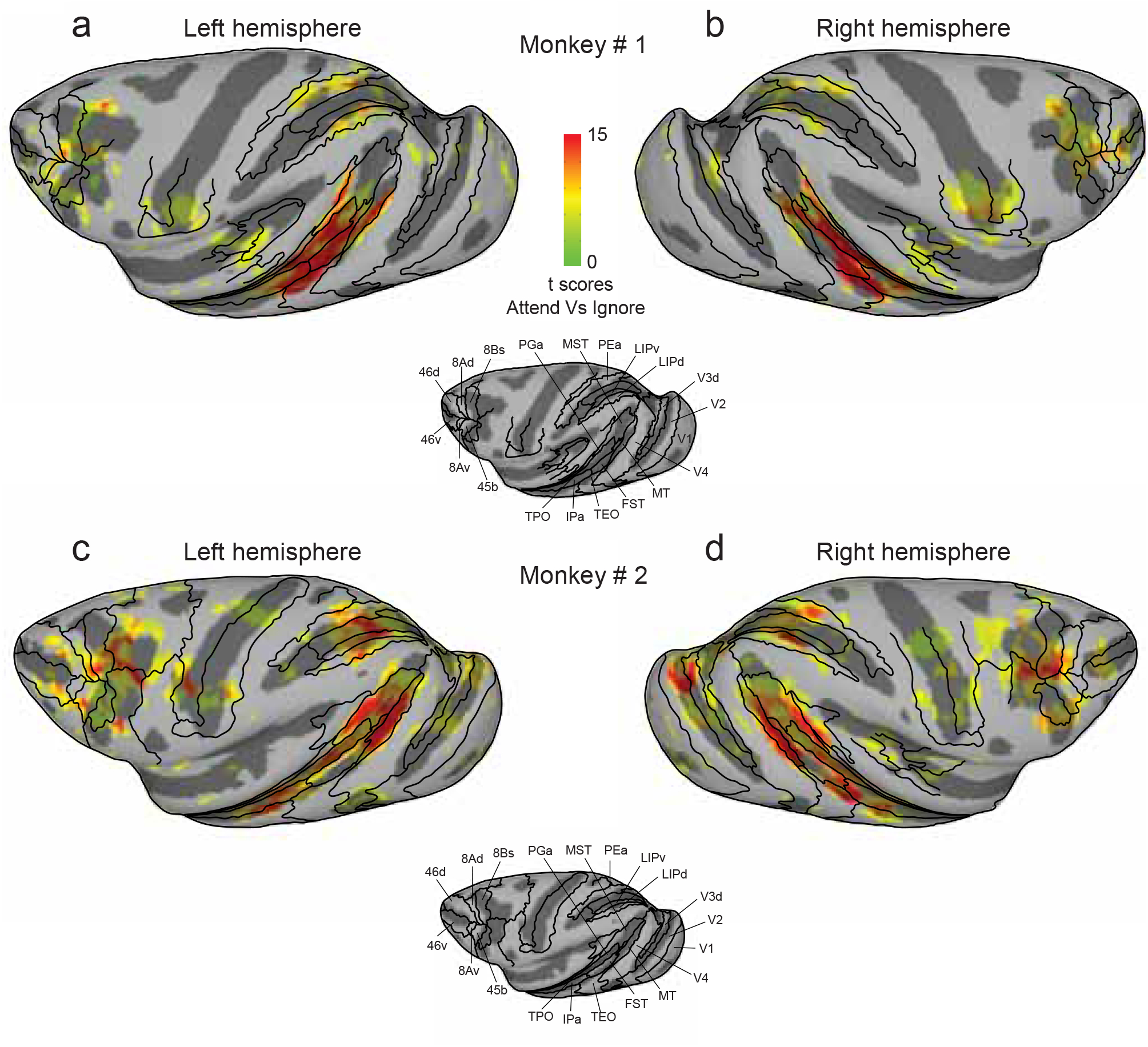
Cortical maps of attention-related activation T-scores contrasting Attend and Ignore tasks were projected onto inflated cortical surfaces of D99 in each monkey’s native space along with anatomical boundaries (black contours). (a, b) Inflated cortical maps of t-scores showing attention-related activation in left (a) and right (b) hemispheres of monkey # 1. Anatomical boundaries are labeled for the left hemisphere in monkey # 1. (c, d) Inflated cortical maps of t-scores showing attention-related activation in left (c) and right (d) hemispheres of monkey # 2. Anatomical boundaries are labeled for the left hemisphere in monkey # 2. T-scores were corrected for multiple comparisons (Bonferroni correction; p < 0.05, t-score > 5.02).

Some of these activated regions were fully expected based on previous studies of visual attention. In posterior STS, we found significant modulation in voxels anatomically identified to be in areas MT and V4t in all four hemispheres (Fig. 3). Monkey #2 showed a larger spread of attention-related modulation in area MST, compared to monkey #1. In the IPS, voxels showing attention-related modulation overlapped with anatomical areas LIPd, LIPv and area PEa (Fig. 3). In the frontal cortex, voxels located in parts of areas 45b, 8Bs, 8Ad in AS (FEF), and area 46v (vlPFC) in PS showed attention-related modulation (Fig. 3).

Unexpectedly, we also found task modulation in different regions of the central sulcus (Fig. 3), but the sites of activation were idiosyncratic across the two monkeys. We speculate that, rather than being due to visual attention, this observed modulation in somatomotor areas was related to differences in how the monkeys used the joystick in the task. Consistent with this interpretation, we did not see any activation in the central sulcus areas during the stimulus mapping experiment in which monkeys passively viewed stimuli during fixation and the joystick was not present (see Supplementary Fig. S1).

The activation maps also consistently revealed a large swath of attention-related modulation in the mid-STS region, anterior to MT and MST. The activated voxels did not fall neatly within a single cortical area, but instead aggregated around the borders of several mid-STS cortical areas (FST, IPa, PGa and TEO) that are more clearly illustrated in a higher magnification view of these same data (Fig. 4). The peak activation in these mid-STS cortical areas was located in the fundus of the STS near the border between areas IPa and anterior FST, but activated voxels extended medially into PGa and TPO onto the dorsal bank of the STS, and laterally into medial parts of area TEO onto the ventral bank. The locations of the peak activation in the fundus varied somewhat across hemispheres, straddling the border between IPa and anterior FST – it was in IPa for the left hemispheres (Fig. 4a, 4c) but in aFST for the right hemispheres (Fig. 4b, 4d) in both monkeys. Based on these results, we will refer to this attention-related region in the fundus of the mid-STS as aFST/IPa.

**Figure 4.**
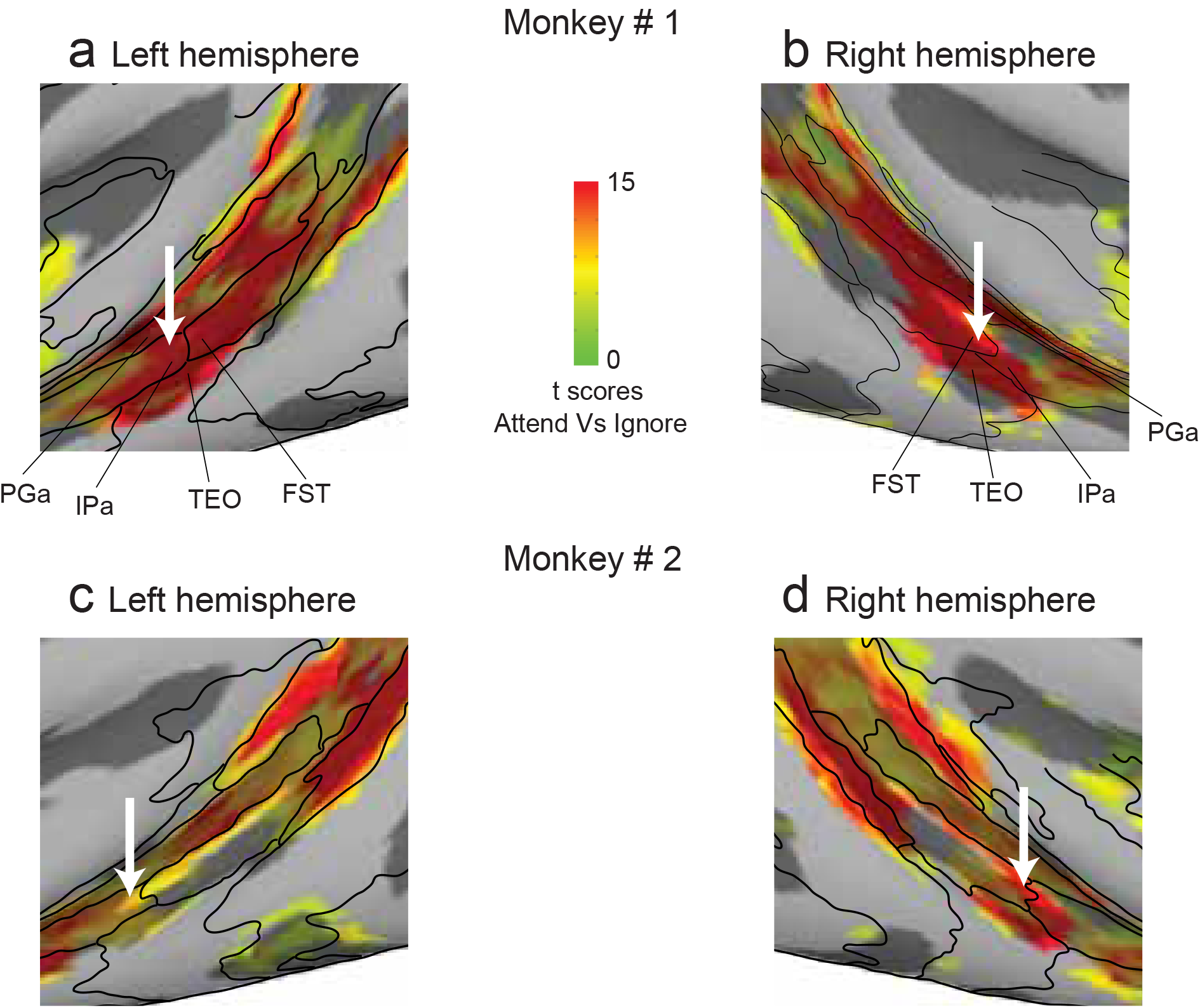
Attention-related modulation in mid-STS areas (a, b) Magnified versions of figure 3a and 3b showing STS activation in left and right hemispheres of monkey # 1. (c, d) Magnified versions of figure 3c and 3d showing STS activation in left and right hemispheres of monkey # 2. The white arrow points to the peak activation in the aFST/IPa region in left and right hemispheres of both monkeys.

To characterize these attention-related modulations in greater detail, we returned to the 3D volumes and identified regions of interest (ROIs) in cortical and subcortical brain regions, defined as 2mm-radius spheres centered on the local maxima of the contrast between Attend and Ignore conditions. For each ROI, we then extracted the time-course of BOLD activity during Attend and Ignore conditions for each area (see Methods). For consistency, we focused on ROIs that showed activation at the same or overlapping anatomical locations in both monkeys. In total, we found 9 cortical and 3 sub-cortical ROIs in both monkeys and assigned each a name based on the anatomical location of the local maxima. The coronal slices and average time-courses of all cortical and sub-cortical attention-related ROIs are shown in Figures 5 and 6, respectively. The average BOLD time-courses for Attend and Ignore conditions in each ROI are displayed next to the corresponding coronal slice. The average BOLD time-course for a given condition in an area was constructed by pooling blocks of that condition across sessions and voxels in that ROI (see Methods).

**Figure 5.**
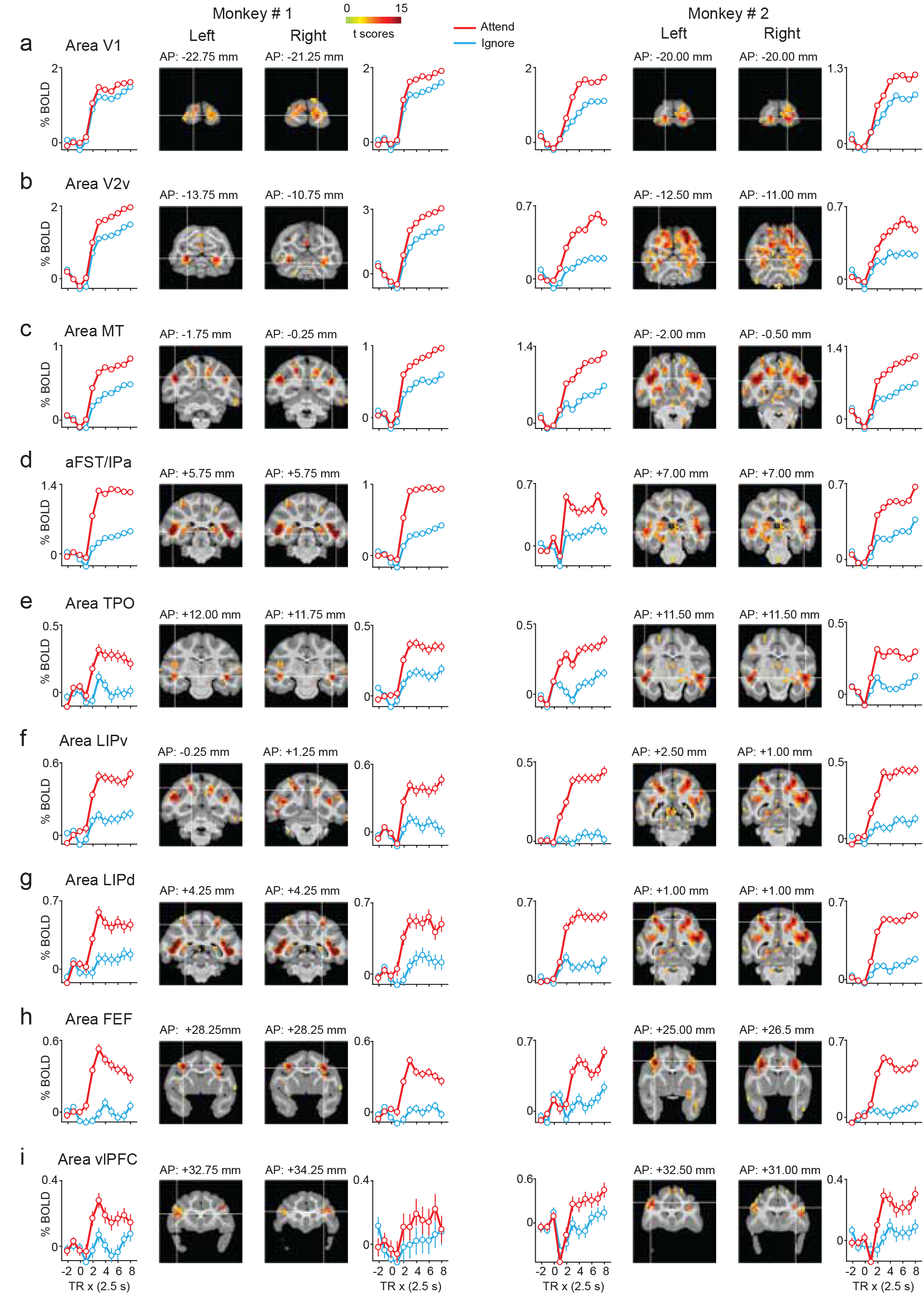
BOLD time-courses during Attend and Ignore tasks in cortical ROIs (a – i) Each row shows plots of mean BOLD time-courses for a given area in left and right hemispheres of both monkeys (Monkey # 1: Left column; Monkey # 2: Right column) along with coronal slices containing the peak of the area. The name of the area is shown at top-left of each row. The location of the coronal slice w.r.t the inter-aural axis is shown on top of each coronal slice. Mean BOLD time-courses are plotted as % change in BOLD on y-axis against repetition time (TR) on the x-axis for Attend (red) and Ignore (blue) tasks. Error bars indicate 95% confidence intervals.TR = 0 on x-axis indicates start of the block. The location of the white cross-hair in each coronal slice indicates the peak of the area in the corresponding hemisphere and is overlaid with attention-related activation (t-scores contrasting Attend and Ignore (Bonferroni correction; p < 0.05, t-score > 5.02), as in figure 3). aFST refers to anterior part of anatomical area FST.

### Attention-related ROIs in the cortex

In agreement with previous electrophysiological studies of covert visual attention in monkeys^1–3,5,7^, we found attention-related peak activations in visual cortical areas including areas V1 and V2v (Fig. 5a, 5b) as well as area MT in posterior STS (Fig. 5c) in all four hemispheres of both monkeys. The visual cortical ROIs (V1, V2v and MT) showed a strong response to visual motion during the Ignore condition as well as during the Attend condition.

In the parietal cortex, attention-related peak activations were observed in areas of the IPS including LIPd and LIPv (Fig. 5f, 5g). In the frontal cortex, we observed attention-related peak activations in areas FEF and vlPFC (Fig. 5h, 5i). These observations are in agreement with the previous neurophysiology studies of attention in monkeys^9–11,14^. Unlike the visual cortical ROIs, the frontal and parietal ROIs (FEF, LIPd, LIPv) showed a strong response to visual motion during Attend but a much weaker response during Ignore (Fig. 5; See also Fig. 2).

Importantly, in the mid-STS cortex, we identified two ROIs based on the attention-related peak activations (Fig. 5d, 5e). One ROI was identified in the dorsal bank of the STS and the peak activation was located in area TPO in all four hemispheres (Fig. 5e). The other ROI was aFST/IPa region (Fig. 5d), which was located in the fundus of the STS. As described earlier, the peak activation of this ROI was always near the border of aFST and IPa, and the 2mm sphere included voxels from several mid-STS cortical areas as defined by the anatomical boundaries: a majority of the activated voxels (62%) belonged to areas aFST and IPa and the remaining voxels (38%) were attributed to areas PGa and TEO. The activations we observed in these two mid-STS ROIs are in the vicinity of STS cortical activations reported in previous monkey fMRI studies^35,36^, a point we consider in more detail in the Discussion.

### *Subcortical attention-related* structures

We also observed significant attention-related modulation in subcortical structures (Fig. 6). In all four hemispheres, we observed significant attention-related modulation in the SC (Fig. 6a) and pulvinar (Fig. 6b), in agreement with previous neurophysiology studies^15,18,19^. In addition to these subcortical regions known to be involved in selective attention, we also observed significant attention-related modulation in the genu of the left caudate nucleus in both monkeys (Fig. 6c). The genu of the caudate receives anatomical projections from motion sensitive areas of STS^40^. A recent fMRI study of covert visual attention in monkeys using static symbols has also reported attention-related modulation in the caudate nucleus, though in its tail rather than its genu^35^.

**Figure 6.**
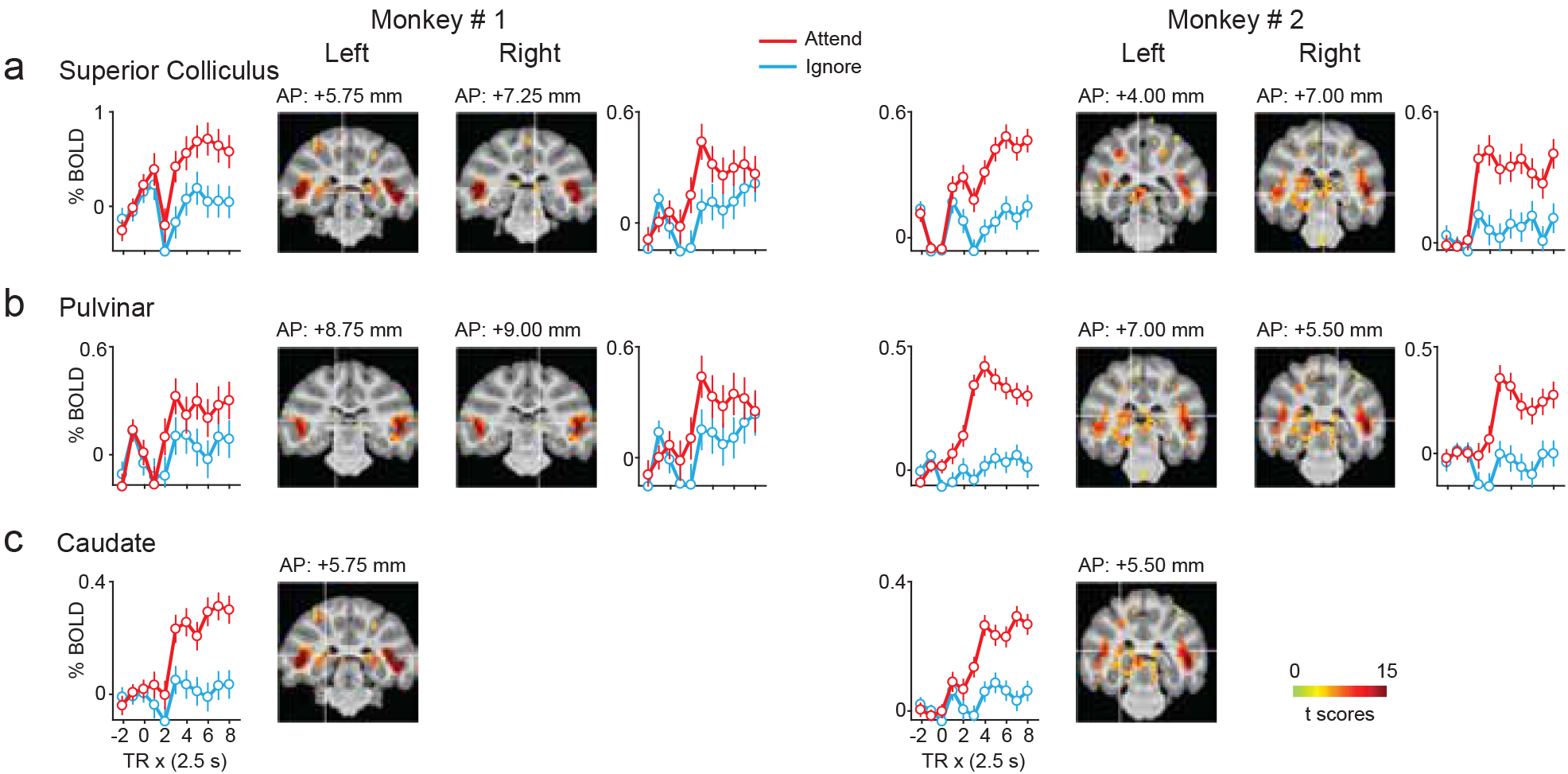
BOLD time-courses during Attend and Ignore tasks in subcortical ROIs (a – c) Same conventions as figure 5.

### Is attention-related modulation in mid-STS regions limited to visual motion stimuli?

A possible explanation for the attention-related modulation in the mid-STS regions (aFST/IPa, TPO) is that these areas are specialized for visual motion processing, similar to areas MT and MST. This explanation would be consistent with previous fMRI experiments showing activations related to higher-order motion signals in similar mid-STS cortical areas^41^. To test how much of the attention-related modulation we observed in these mid-STS regions was due to our use of a visual motion stimulus, we collected data from a control experiment in one monkey (monkey # 1) performing an attention task in the scanner involving a second-order orientation detection (see Methods). All aspects of the task were the same as in the motion version except that the motion stimulus was replaced by a second-order orientation stimulus. Performance of the monkey in the three tasks (Baseline, Ignore and Attend) is shown in Figure 7. The hit and false alarm rates in the Baseline condition were 73% and 10% respectively (Fig. 7b). The hit rate (72%) and foil false alarm rate (Left: 12%, Right: 9%) in the Ignore condition show that the monkey successfully ignored the peripheral second-order orientation stimuli when they were irrelevant (Fig. 7c). The hit rates (Left: 79%, Right: 65%) of the monkey in the Attend condition were higher than false alarms (8%), indicating that the monkey was able to detect the second-order orientation stimulus when it was behaviorally relevant during the Attend blocks (Fig. 7d).

**Figure 7.**
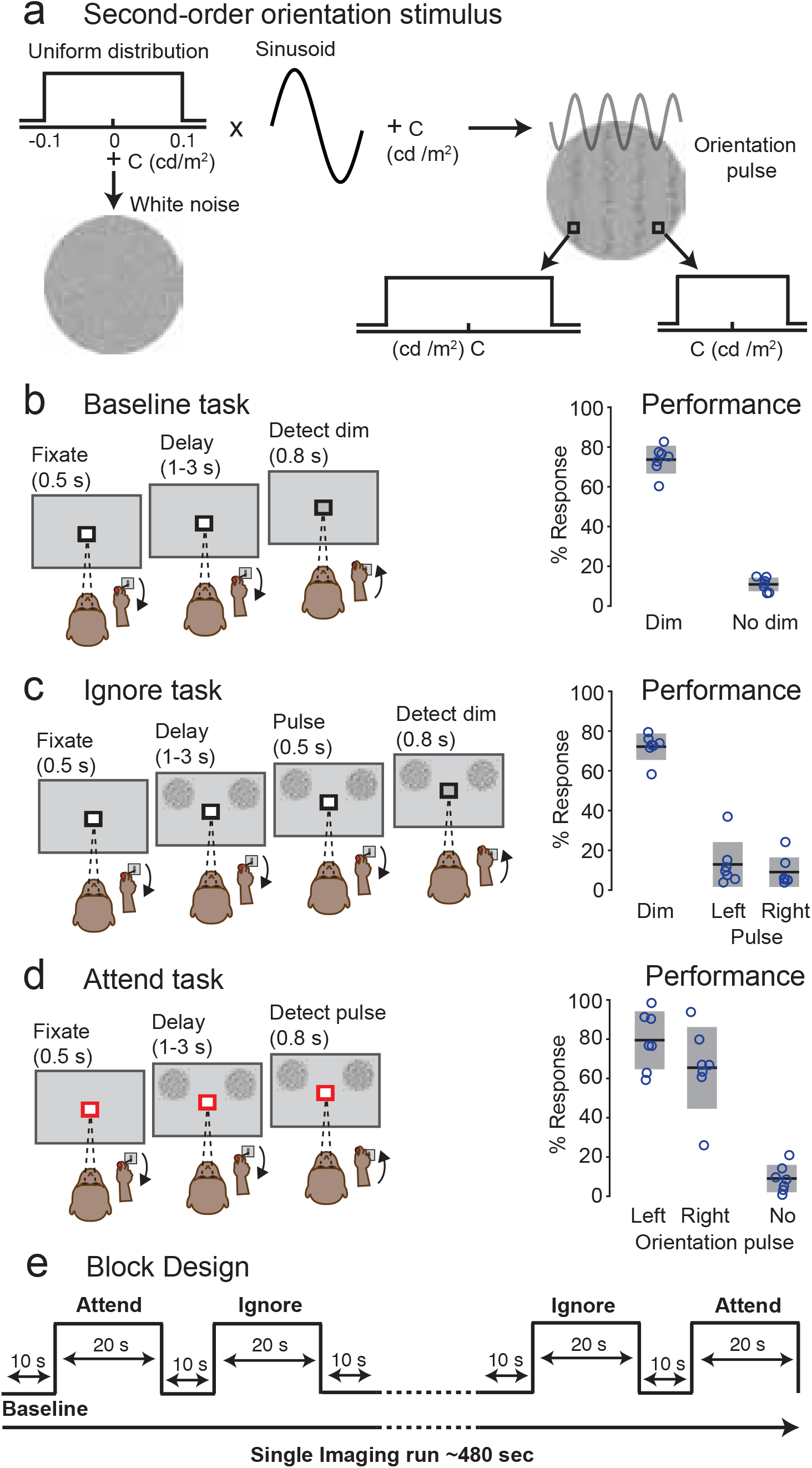
Second-order orientation stimulus: behavioral tasks and performance a) Second-order orientation stimulus. A 2D uniform distribution centered at zero was added to a base luminance (c) to generate a white noise stimulus. The same 2D uniform distribution was modulated by a 2D sinusoid and added to a base luminance to generate a second-order orientation stimulus. The orientation seen in the stimulus is second-order, because it is based on the local contrast of the bands of the grating without any difference in the local mean luminance of the bands of the grating. (b – d) All task conventions were the same as in figure 1, except that the peripheral stimuli were second-order orientation patches rather than visual motion patches. e) Block Design: All three tasks were presented in a block design identical to that used in the motion version of the task (Figure 1).

As with the motion task, we contrasted the Attend and Ignore conditions to identify voxels with significant attention-related modulation. The resulting t-score maps (Bonferroni corrected; p<0.05, t-scores > 5.02) projected onto partially inflated cortical surfaces for monkey #1 are shown in Figures 8ab. The overall attention-related modulation was sparser during the second-order orientation task than the motion-change detection task; early visual areas V1, V2v did not show any significant attention-related modulation. Voxels showing attention-related modulation were identified in areas neighboring the superior temporal sulcus (STS), intra-parietal sulcus (IPS) and arcuate sulcus (AS). In the posterior STS, attention-related modulation was very weak and included only area V4t but not MT (Fig. 8a, 8b), unlike in the motion-change detection task which included both MT and V4t (Fig. 8a, 8b). In the IPS, significantly modulated voxels were located in areas LIPd, LIPv and area PEa. In the AS, voxels were identified in areas 8Bs and 8Ad (FEF). In the mid-STS, significant attention-related modulation was found in the aFST/IPa region with activated voxels identified to be in anterior FST, and IPa in the fundus of the STS, area PGa in the dorsal bank and medial parts of area TEO in the ventral bank. However, we did not find any activation in area TPO on the dorsal bank. These results show that the aFST/IPa region was modulated during covert visual attention in the absence of any visual motion.

**Figure 8.**
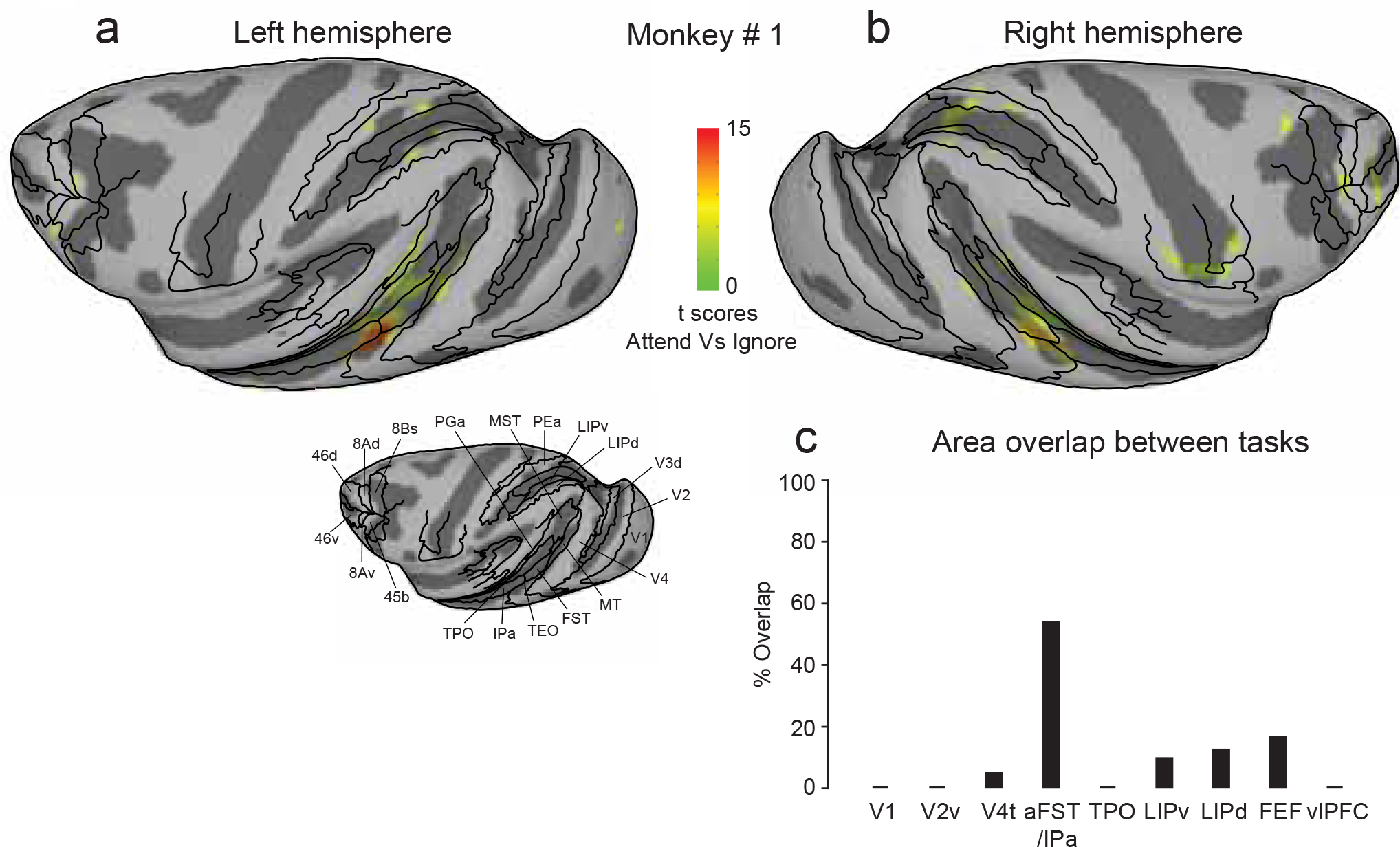
Cortical areas showing attention-related modulation to second-order orientation stimulus (a, b) T-scores contrasting Attend and Ignore tasks described in figure 7 were projected onto inflated cortical surfaces of D99 in native space of monkey # 1 along with anatomical boundaries (black contours). Inflated cortical maps of t-scores (Bonferroni correction; p < 0.05, t-score > 5.02) showing attention-related activation in left (a) and right (b) hemispheres of monkey # 1. Anatomical boundaries for the left hemisphere of monkey # 1 are shown in figure 3a. c) % Overlap for a given area was defined as the percentage of total voxels in that area that showed significant attention-related modulation in both motion (figure 1) and second-order orientation (figure 7) versions of the tasks.

To test if the modulated voxels identified in the motion-change detection and orientation detection tasks were from the same or different population of voxels, we identified attention-related ROIs areas in the orientation task following the same method as previously described for the motion task. Note that the ROIs were defined independently for data from each task based on their respective attention-related modulation. We then computed the percentage of voxels for each ROI that were significantly modulated during both attention tasks. We defined the “%Overlap” for an area as the number of voxels that showed significant attention-related modulation in both tasks, divided by the total number of voxels in the joint ROI (i.e., the union of the voxels across the two tasks). A low value for %Overlap would indicate that the attention-related modulation depended on the particular visual feature (motion or orientation) used in the attention task, or that the center of the ROI shifted considerably between the two tasks; a high value would indicate that the same voxels were modulated in both tasks. We found modest values for %Overlap for frontal and parietal ROIs (LIPd: 9%, LIPv: 12%, FEF: 16%), and a relatively low value for the V4t ROI (4%) neighboring MT (Fig. 8c). The highest value for %Overlap was found for aFST/IPa (53%), demonstrating that the attention-related modulation in the aFST/IPa region was not limited to attention tasks involving visual motion – indeed, nearly half of the voxels in the aFST/IPa ROI were modulated in both tasks (Fig. 8c).

As an additional comparison of the aFST/IPa modulation during the two attention tasks, we examined the full extent of the overlap using the activation maps for the entire mid-STS region, rather than considering only the voxels within the 2mm-radius sphere around the peak activation. The blue contour in Fig. 9a, 9b outlines the contiguous attention-related activation in the mid-STS region during the orientation task for both hemispheres in monkey #1, which extended across anterior FST, IPa and PGa. We then overlaid the same blue contour on the activations during the motion task, and found that the highest activations during the motion task were mostly contained within the blue contour (Fig. 9c, 9d). This demonstrates that not only was the aFST/IPa region modulated during both tasks, but the voxels modulated during the orientation task were the same voxels that showed the largest modulation during the visual motion task.

**Figure 9.**
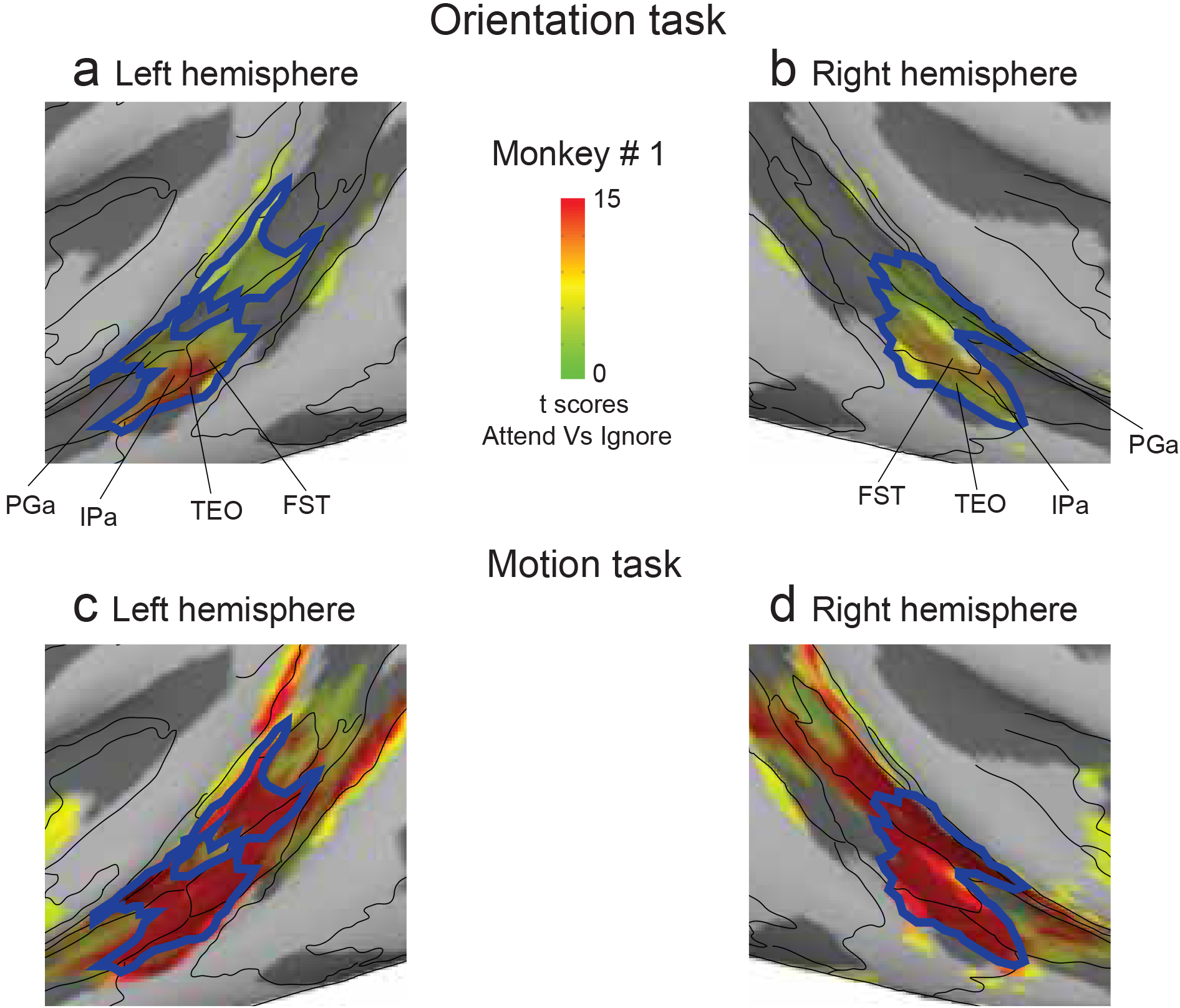
Overlap of attention-related modulation in mid-STS areas during orientation and motion tasks (a, b) Magnified versions of figure 8a and 8b showing STS activation during orientation task in left and right hemispheres. Blue contour represents the contiguous activation in mid-STS areas. (c, d) Magnified versions of figure 3a and 3b showing STS activation during motion task in left and right hemispheres. Blue contours from figure 9a and 9b were overlaid for the left and right hemispheres respectively.

Finally, we considered whether the difference in the spatial extent of the attention-related activations between the motion and orientation tasks (t-score maps in Fig. 3 and Fig. 8) might be due to the difference in the number of trials of imaging data in Motion task (5244 Attend trials) and Orientation task (3602 Attend trials). We repeated the GLM analysis (See Methods) on an independent dataset collected during the motion task with a number of trials (3667 Attend trials) comparable to those in the orientation task. The resulting attention-related activation map showed significant activations around the superior temporal sulcus, intra-parietal sulcus, arcuate sulcus and principal sulcus, comparable to the attention-related activations shown in Fig 3. These results indicate that the difference in attention-related activations between the two tasks was not due to the difference in the number of runs or trials of imaging data collected, but instead due to the difference in the visual stimuli used. In particular, we suspect that the dynamic nature of the visual motion stimulus simply produced larger activations overall than the second-order orientation stimulus.

## Discussion

The goal of the present study was to replicate the results from previous monkey fMRI studies in a different attention task paradigm and identify the brain regions modulated during covert visual attention in non-human primates. The task was directly comparable to that used in a previous human fMRI study to identify the brain areas that show attention-related modulation to visual motion^39^. We found attention-related modulation in visual cortical areas (V1, V2v, MT), fronto-parietal areas (FEF, vlPFC, LIP) and sub-cortical structures (SC, pulvinar), as might be expected from previous electrophysiology studies in monkeys^1–3,5,8,9,12–15,18,19^. These observations are in agreement with the recent fMRI studies of covert visual attention in monkeys^34–38^, as well as the more numerous fMRI studies of covert visual attention in humans^39,42–44^. Interestingly, our results also show attention-related modulation in relatively unexplored areas of the brain – mid-STS areas (aFST/IPa region and area TPO) and caudate nucleus during covert visual attention to motion in non-human primates (Fig. 3, 4 and 5).

Areas in mid-STS (aFST/IPa region and area TPO) showed modulation during covert attention to motion. The anatomical location of the aFST/IPa region (Fig. 4d; Monkey #1: AP + 5.75mm; Monkey # 2: AP + 7mm) matches the previously described motion-sensitive region in STS named LST^41^. To test if the attention-related modulation we observed in aFST/IPa region was limited to visual motion stimulus, we used the same task design with a second-order orientation stimulus, rather than a motion stimulus. Areas of the fronto-parietal network (FEF, LIP) showed some modulation in both versions of the attention task, as might be expected, but we also found strong modulation in aFST/IPa region located in the mid-STS (Fig. 8a, 8b). In fact, we found that modulation in aFST/IPa region was localized to the same voxels in both tasks (Fig. 8c), according to the D99 atlas in AFNI^45^. The LST region was also shown to exhibit a preference for intact shapes over scrambled shapes^41^, consistent with our observation that attention-related modulation in this region was not restricted to tasks using visual motion (Fig. 8a, 8b). Although the LST region was not previously shown to be modulated by attention, our results suggest that aFST/IPa region is the previously described LST, and this region is recruited during covert visual attention even without visual motion. The functional contribution of these mid-STS areas (aFST/IPa region and area TPO) to covert visual attention is not known, although there is sparse evidence that lesions targeted in the fundus and dorsal bank of STS can produce unilateral neglect in monkeys^46,47^.

We also found strong attention-related modulation in the genu of the caudate nucleus during covert attention to motion. Previous fMRI studies in monkeys using static symbols found modulation in the tail of the caudate nucleus^35^. This difference is most likely due to difference in the visual stimuli used: visual areas involved in processing the visual motion stimuli we used in our task project to caudate genu, whereas visual areas involved in processing object stimuli used by Caspari et al. project to caudate tail^40^. Furthermore, lesions of the striatum including caudate nucleus leads to hemi-spatial neglect in humans^48^. Our results taken together with the observations from Caspari et al. suggest that the caudate nucleus in monkeys plays some role in the performance of covert attention tasks. Neuronal recordings in the caudate have revealed activity related to perceptual and cognitive functions^49^, including the encoding of object values^50^ and the formation of perceptual choices based on visual motion signals^51^, but the activity of caudate neurons has not yet been reported for covert visual attention tasks. Circuits involving the caudate nucleus might contribute to covert visual attention through mechanisms related to the formation of the perceptual choice^52,53^, rather than the modulation of sensory processing, but additional studies will be needed to sort out these issues.

The attention-related modulation in cortical and sub-cortical areas we observed is broadly consistent with the results of previous fMRI studies in monkeys during covert attention tasks^34–38^. One recent study investigated attention-related modulation in monkeys using visual motion stimulus, but in a different task paradigm^36^. The results from Stemmann et al. showed that an area in posterior inferotemporal cortex exhibited a strong attention-related modulation during covert attention to motion^36^. The majority of the attention-related activation we found was in the fundus and dorsal bank of the STS, including area TPO and a region we referred to as the aFST/IPa region. We also observed some attention-related activation in the medial parts of area TEO in the lower bank of the STS. The anatomical location of the activation in the Stemman (2016) study (see their Fig. 2D) is at least 6mm posterior to the activations we observed (our Fig. 4d), even though the relative location of these STS activations with respect to MT is the same in both studies (compare Fig. 2D in Stemmann et al. and Fig. 4 in our study). We thus suspect that the anterior-posterior locations of our STS activations might be very similar. The discrepancy in medial-lateral location of activations in both studies might be explained by the difference in the retinotopic location of the stimuli used (8^0^ in our study compared to 5^0^ used in Stemmann et al.), given the retinotopic organization in mid-STS areas^54,55^. Regardless of these issues, our results taken together with Stemmann et al. provide strong evidence that the aFST/IPa region is not only modulated during attention to motion stimuli but also during attention to other visual features.

In conclusion, using fMRI in two monkeys performing a covert attention task, we identified a list of brain structures that are selectively activated during covert attention to peripheral visual stimuli. In addition to attention-related activation in expected cortical (frontal, parietal and visual areas) and subcortical (superior colliculus, pulvinar) regions, we also found significant attention-related modulation in places not traditionally linked to attention – mid-STS areas (aFST/IPa region and area TPO) and the caudate nucleus. These findings identify the mid-STS areas and caudate as additional brain areas of interest for the study of covert visual attention in non-human primates.

## Methods

### Animals

Two adult male rhesus monkeys (*Macaca mulatta*) weighing 7-9 kg participated in this study. All experimental protocols were approved by the National Eye Institute Animal Care and Use Committee and all procedures were performed in accordance with the United States Public Health Service policy on the humane care and use of laboratory animals.

### Attention tasks: Motion-change detection

In the scanner, both monkeys performed three behavioral tasks: Baseline, Ignore and Attend. In all tasks, monkeys initiated the trial by holding the joystick down and fixating the central fixation spot with a colored central cue on a grey background. Monkeys fixated for the entire duration of the trial with in a 2° fixation window. The color of the central cue indicated the trial condition. In Baseline and Ignore trials, the color of the central cue was black (Fig. 1a, 1c) and the relevant stimulus was the fixation stimulus. In Attend trials, the color of the central cue was red (Fig. 1e) and the relevant stimulus was the peripheral motion stimulus. The details about the peripheral motion stimuli and fixation stimuli are provided elsewhere (see Supplementary methods). The sequence of events in three different trial conditions are as follows.

In Baseline trials, following 0.5 s of fixation, the luminance of the fixation spot decreased during a variable delay of 1 – 3 s on the 65% of the trials. Monkeys reported the luminance change by releasing the joystick within 0.3 - 0.8 s to get a juice reward (Fig. 1a). A total of 12340 Baseline trails were collected in both monkeys (7520 in Monkey #1 and 4820 in Monkey #2).

In Ignore trials, following 0.5 s of fixation, two random dot motion stimuli were presented on either side of fixation at 8° eccentricity (radius) and 10° above horizontal meridian (azimuth). During the variable delay of 1 – 3 s, the luminance of the fixation spot decreased on 65% of the trials. Independent of the fixation luminance change, one of the peripheral motion stimulus changed direction during the variable delay of 1 – 3 s on the 65% of the trials. Monkeys ignored the motion direction change and reported the luminance change by releasing the joystick within 0.3 - 0.8 s to get a juice reward (Fig. 1c). If the monkeys released the joystick for a motion direction change, the trial was aborted. A total of 13189 Ignore trails were collected in both monkeys (7927 in Monkey #1 and 5262 in Monkey #2).

In Attend trials, following 0.5 s of fixation, two random dot motion stimuli were presented at the same stimulus location as in Ignore trials. One of the peripheral motion stimulus changed direction during the variable delay of 1 – 3 s on the 65% of the trials. Monkeys reported the motion-direction change by releasing the joystick within 0.3 - 0.8 s to get a juice reward (Fig. 1e). There was no fixation luminance change in Attend trials. A total of 12424 Attend trails were collected in both monkeys (7180 in Monkey #1 and 5244 in Monkey #2).

In all tasks, 35% of the trials were catch trials and monkeys hold the joystick down to get a juice reward.

### Attention tasks: Orientation-pulse detection

In addition to the motion-change detection task, monkey # 1 also performed a version of the attention tasks with orientation pulse stimuli instead of the random dot motion stimuli. The sequence of events in all three conditions (Baseline, Ignore, Attend) were kept the same as the motion-change detection version of the task (Fig. 7b, 7c, 7d). The onset of the motion stimuli was replaced with the onset of white noise stimuli, and the motion-direction change was replaced with a 0.5 s second-order orientation pulse. In Attend condition, monkey reported the orientation pulse by releasing the joystick within 0.3 - 0.8 s to get a juice reward (Fig. 7d), whereas in Ignore condition, monkey ignored the orientation pulse and reported the luminance change in the fixation spot by releasing the joystick within 0.3 - 0.8 s to get a juice reward (Fig. 7c). A total of 3518 Ignore trails and 3602 Attend trials were collected in in Monkey #1.

The second-order orientation stimulus was generated by briefly (0.5 s) modulating the contrast of a white noise stimulus with a 2-dimensional sinusoid (Fig. 7a). The noise stimulus was 6^0^ in diameter and consisted of checks each the size of a pixel with luminance values ranging from 8 – 84 cd/m^2^, and the 2-dimensional sinusoid had a spatial frequency of 0.7 cycles/deg, and its orientation was 90^0^. We refer to this as a second-order orientation stimulus, because the oriented grating briefly visible in the stimulus was due to the local differences in contrast, not luminance differences. The mean luminance (38 cd/m^2^) of the stimulus was constant throughout its presentation and was the same across every band in the oriented grating.

### Block Design

Baseline, Ignore and Attend tasks were presented in a block design as shown in Fig. 1g. Each run started with Baseline block which lasted for 10 s and was presented in every alternate block thereafter. Ignore and Attend blocks lasted for 20 s and were presented randomly between Baseline blocks. The number of Ignore and Attend blocks were balanced in a given run. Each run lasted 480s. For the motion-change detection task, a total of 324 runs (132 in Monkey #1; 192 in Monkey # 2) were collected in both monkeys across 24 sessions. For the orientation-pulse detection task, a total of 102 runs were collected in monkey # 1 across 7 sessions.

### fMRI data collection

Anatomical and functional images were collected in a vertical magnet (4.7T, 60 cm vertical bore; Bruker Biospec) equipped with a Bruker S380 gradient coil. EPI volumes were acquired using a custom built transmit and 4-channel receive RF coil system (Rapid MR International). In each run, we collected 192 whole-brain EPI volumes at an isotropic voxel resolution of 1.5 mm and at a TR of 2.5 s.

### fMRI Analysis

Preprocessing of the fMRI data was done using AFNI/SUMA software package^56^. Raw images were converted to AFNI data format. EPI volumes in each run were slice-time corrected using *3dTshift*, followed by correction for static magnetic field inhomogeneities using the *PLACE* algorithm^57^. All EPI volumes were motion-corrected using *3dvolreg* and were registered to the session anatomical using *3dAllineate*. To combine EPI data across multiple sessions for a given animal, all sessions for a given animal were registered to a template session. A high resolution anatomical of each monkey was registered to the corresponding template session anatomical to overlay functional results in monkey’s native space. To overlay D99 atlas boundaries on the functional results, D99 anatomical in AFNI was registered to each monkey’s native space^45^. Surface maps were generated for D99 anatomical in each monkey’s native space using CARET from a white matter segmentation mask^58^. Surface maps of D99 in each monkey’s native space were viewed in SUMA overlaid with anatomical boundaries and functional results.

### Functional maps of attention

To identify voxels that were modulated by attention we performed a GLM analysis using *3dDeconvolve* in AFNI. Attend and Ignore conditions were included as regressors of interest and Baseline condition, motion correction parameters were included as regressors of no interest (baseline model). To control for the effects caused by any differences during Attend and Ignore blocks in blinks (Wilcoxon rank sum test, p = 0.91 (Monkey # 1), p = 0.97 (Monkey # 2)), saccadic eye movements (Wilcoxon rank sum test, p = 0.04 (Monkey # 1), p < 0.0001 (Monkey # 2)), rewards (Wilcoxon rank sum test, p = 0.17 (Monkey # 1), p = 0.68 (Monkey # 2)) and joystick movements (Wilcoxon rank sum test, p = 0.11 (Monkey # 1), p < 0.001 (Monkey # 2)), we included these factors as part of the regression model. Reward times, joystick event times (press and release), blink times and saccade times (left and right) were convolved with the hemodynamic impulse response function and were included as part of the baseline model. The median duration of fixation in the blocks of all three conditions for both monkeys is above 85%. Saccades with magnitude less than 1° were detected using velocity and acceleration threshold^59^. T-scores contrasting Attend and Ignore conditions were projected on to the inflated maps to show voxels modulated by attention. All functional maps were thresholded (p < 0.05; t-score > 5.02) to correct for multiple comparisons (Bonferroni correction).

### Attention-related areas and BOLD time-courses

Attention-related areas were identified based on the local maxima of the attention activation map for each hemisphere using *3dExtrema* in AFNI. The BOLD time-course for each Attend or Ignore block was computed as a % change in BOLD relative to the BOLD in Baseline block preceding it. For each area, an average time-course was constructed by pooling time-courses of all voxels within a 2mm radius around the local maxima across all blocks from all sessions.

## Data Availability

The datasets generated during and/or analysed during the current study are available from the corresponding author on reasonable request.

## Acknowledgments

We thank Tom Ruffner and Nick Nichols for technical support and Fabrice Arcizet, James Herman, Leor Katz and Lupeng Wang for helpful input. We also thank Brian Russ for helpful suggestions in setting up fMRI preprocessing pipeline and constructing inflated maps. This work was supported by the National Eye Institute Intramural Research Program at the National Institutes of Health. Functional and anatomical MRI scanning was carried out in the Neurophysiology Imaging Facility Core (NIMH, NINDS, NEI) supported under IRP grant ZIC MH002899. This work utilized the computational resources of the NIH HPC Biowulf cluster.

## Author Contributions

All authors designed the experiments. A.R.B. and A.B. conducted experiments and collected the data. A.R.B. analyzed the data. All authors interpreted the results. A.R.B. and R.J.K wrote the manuscript.

